# Protein kinase A gene knockout in *Coprinopsis cinerea* by CRISPR/Cas9 RNP complex system

**DOI:** 10.1101/2022.10.28.513995

**Authors:** Po Lam Chan, Hoi Shan Kwan, JinHui Chang

**Affiliations:** School of Life Sciences, The Chinese University of Hong Kong, Shatin, New Territories, Hong Kong SAR, China; Research Institute for Future Food, The Hong Kong Polytechnic University, Hong Kong SAR, China

## Abstract

*Coprinopsis cinerea* is a model mushroom for studying the developmental processes of homobasidiomycetous fungi. The development of *C. cinerea* depends on the sensing of environmental conditions, such as light, temperature, humidity and nutrients. The signal transduction pathway from the environmental condition sensors to the regulators and reproduction is an important research topic as the knowledge on it is still fragmentary. Protein kinases not only are crucial to the signal transduction, but also can be master switches of development. However, information on their role in basidiomycetous fungi is still limited. Using the effective gene-targeting of the current gene modification, a system of CRISPR/ Cas9 RNP with double strain template donor that contained a selective mark hygromycin cassette was developed in this study. In this system, *CcPka1* gene knockout was successfully generated through homologues repair together with Cas9 RNP complexes. This was the first report to use CRISPR/ Cas9 RNP approach with C. cinerea. Δ*CcPka1* mutants showed slow mycelial growth and rapid fruiting body induction compared to the wild type even when under high glucose condition. In addition, Δ*CcPka1* mutants had significantly increased intracellular glucose level and decreased glycogen level. Moreover, Δ*CcPka1* mutants regulated the gene expression of components in cAMP-PKA pathway. All these results demonstrated that PKA played an important role in the nutrient signal transduction and negatively regulated the fruiting body formation. This study also confirmed that GSK3 could be the downstream target of PKA, but their interaction requires further investigation.

## Introduction

Fungi can sense and respond to in environmental changes. Nutrient depletion is a critical parameter of worsening environment and a crucial signal for sexual reproduction of fungi with fruiting body formation (Aschan-Äberg 1958; Plunkett 1953). The culturing of the fungus *Coprinus macrorhizus* with high glucose substrates inhibited the formation of fruiting body (Uno and Ishikawa 1974). The glucose-induced signal coordinates with different signaling pathways (Rolland et al. 2000). cAMP-dependent protein kinase A (PKA) signaling pathway is a crucial nutrient-sensing pathway. Adenylyl cyclase in this pathway converts ATP to cAMP, leading to the activation of PKA by binding to the regulatory subunit of PKA to release the catalytic subunit of PKA (Sands and Palmer 2008). This signaling pathway regulates a variety of cellular functions through activation or repression of its downstream targets. For example, in *Saccharomyces cerevisiae*, the PKA pathway mediates glucose uptake and regulates carbohydrate metabolism (Freitas et al, 2010). The PKA pathway in *Cryptococcus neoformans* regulates mating and virulence (Hicks et al. 2004). The overexpression of PKA in *Schizophyllum commune* suppresses vegetative growth and fruiting body formation (Yamagishi et al. 2005). There are few studies on the role of the PKA pathway in sexual reproduction in mushroom-forming fungi.

To investigate the role of the cAMP-dependent PKA pathway in nutrient-sensing and fruiting body formation of *Coprinopsis cinerea* (*C. cinerea*), a target gene modification method, CRISPR/Cas9 RNP, was developed to knock out the *C. cinerea* PKA gene (*CcPka1*, CC1G_01089) that encodes a PKA catalytic subunit. This is the first time CRISPR/Cas9 RNP has been used to generate mutants in *C. cinerea*. This study showed PKA transduced nutrient signals and induced fruiting body formation.

In addition, the results proposed that PKA and Glycogen Synthase Kinase (GSK3) interacted to regulate intracellular glycogen synthesis during vegetative growth and fruiting body formation in *C. cinerea*.

## Materials and methods

### Strain and culture medium

*C. cinerea* strain # 326, a homokaryotic fruiting strain, was grown in a yeast extract-malt extract-glucose (YMG) medium solidified with 1.5% (w/v) agar. Mycelia were harvested for genomic DNA extraction using the DNeasy Plant Mini Kit (Qiagen). For fungal transformation, the regeneration medium supplemented with an antibiotic, Hygromycin B (Sigma-Aldrich), was used to select the potentially mutated candidates.

### Design and selection of sgRNA for target gene

The sgRNA was designed by the bioinformatics tool sgRNAcas9 (Xie et al. 2014). Parameters for candidate protospacers were as follows: the sgRNA binding site was located close to the 5’ end of the target gene coding region, the protospacer and PAM sequence was (N)20-NGG, and the number of mismatches of the protospacer was as few as possible when mapping to the whole reference genome of *C. cinerea*. Two sgRANs were selected from *CcPka1* (CC1G_01089), encoding as PKA catalytic subunit, based on the fewest off-target site criterion. They were located on 34 bp and 48bp on the exon 2 and exon 3 of the coding region, respectively. The selected sgRNAs are shown in Table 1. The sgRNAs were ordered as Alt-R CRISPR-Cas9 cr.

**Table 1.**
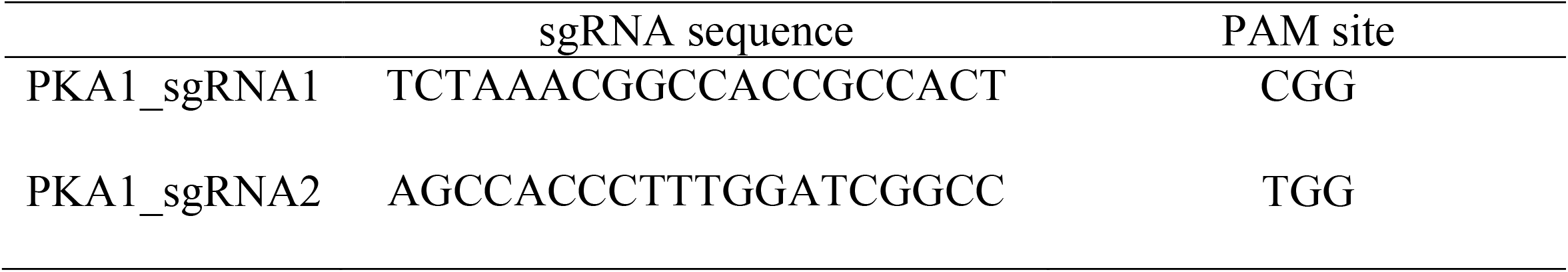
Selected sgRNAs for CRISPR-Cas9 knock out of target gene.

### Plasmid construction for homologous repair template donor

The homologous repair template donor was constructed by using overlap extension PCR with two rounds of PCR. In the first round of PCR, 280 bp upstream homologous flanking sequence and 295 bp downstream homologous flanking sequence from sgRNA1 cut site and sgRNA2 cut site in the *CcPka1* gene were amplified from the *C. cinerea* genomic DNA using the KAPA HiFi HotStart PCR kit (Kapa Biosystems, Wilmington, MA, USA). The PCR thermocycling profile used was as follows: initiation denaturation for 3 minutes at 95°C, followed by 30 cycles of 20 seconds at 98°C, 15 seconds at 60°C, and 30 seconds at 72°C, followed by a final extension for 1 minute at 72°C and held at 4°C. The hygromycin cassette including the *B-tubulin* promoter and terminator was amplified from pPHT1 plasmid with the KAPA HiFi HotStart PCR kit (Kapa Biosystems, Wilmington, MA, USA). The PCR thermocycling profile used was as follows: initiation denaturation for 3 minutes at 95°C, followed by 30 cycles of 20 seconds at 98°C, 15 seconds at 60°C, and 1minute and 45 seconds at 72°C, which was followed by a final extension for 1 minute at 72°C and held at 4°C. Three PCR amplicons in the first round of PCR were visualized on 1% agarose gel with SYBR Safe DNA gel stain (Invitrogen). Then the PCR amplicons were cut from the gel and purified with Gel/PCR DNA Fragment Extraction Kit (Geneaid Biotech, Taiwan).

In the second round of PCR, 3 purified PCR fragments acted as templates were amplified with primers containing restriction enzyme cut site BamHI and XhoI in the upstream homologous flanking region and downstream homologous flanking region. The PCR thermocycling profile used was as follows: initiation denaturation for 3 minutes at 95°C, followed by 35 cycles of 20 seconds at 98°C, 15 seconds at 60°C, and 2 mintues and 30 seconds at 72°C, which was followed by a final extension for 1 minute at 72°C and held at 4^***o***^C. The PCR amplicons were visualized on 1% agarose gel with SYBR Safe DNA gel stain (Invitrogen) and purified with Gel/PCR DNA Fragment Extraction Kit (Geneaid Biotech, Taiwan). The homologous repair template donors were inserted into the BamHI and XhoI enzyme site of pcDNA3. The assembled plasmids were sequenced to confirm they were correct and were extracted with DNA-spin Plasmid DNA Purification Kit (iNtRON Biotechnology). All the primers used for template donor and plasmid construction and sequencing in this study are listed in Table 2.

**Table 2.**
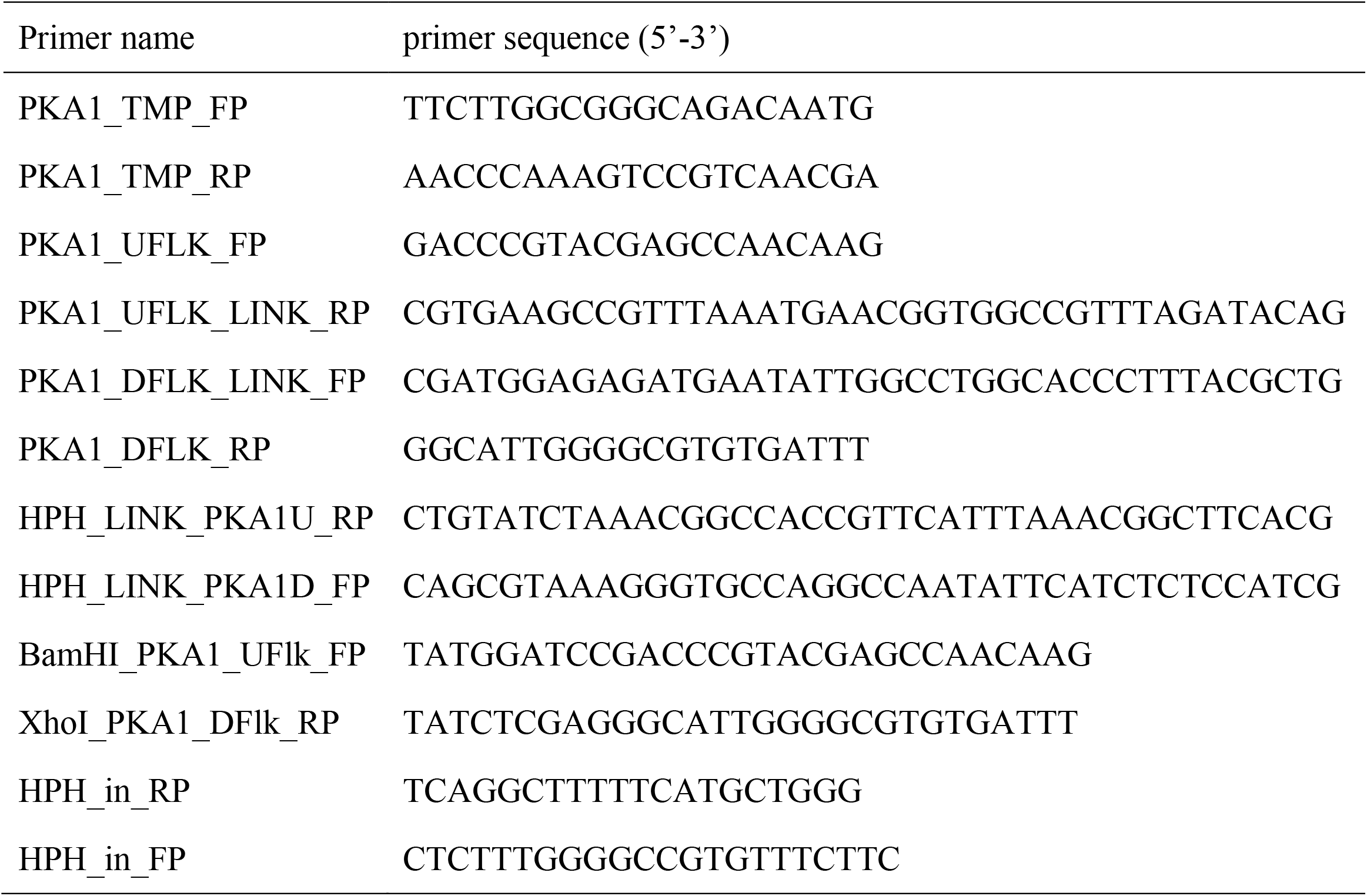
Primers for plasmid construction and mutant verifying by PCR.

### PEG-mediated fungal transformation

*C. cinerea* was grown in a 60mm petri dish at 37°C with light for 6 days to generate oidia. To harvest the oidia, 5ml distilled water was added to the surface of the petri dish and the surface of *C. cinerea* was brushed gently. The oidia solution was collected in a 15ml falcon tube and centrifuged at 2.600xg for 5 mins. The supernatant was discarded and resuspended in 5ml sterile MM buffer. The solution was centrifuged at 2,600xg for 5 mins and the supernatant was discarded. The oidia pellet was then digested with 10mg/ml Novozyme (Novo BioLabs, Denmark) at 37°C with shaking at 110rpm for 3-4 hours. 10µl of the mixture was taken for checking the time interval of the digestion of the oidia. When the rod shape of the oidia became round, MMC buffer was added to stop the digestion of Novozyme and protoplasts were washed twice. Around 0.2-2×10^8 protoplasts were collected in 500µl MMC buffer.

Before the PEG-mediated transformation, 6µl 25µM sgRNA1 and sgRNA2 were mixed with 2µl Alt-R S.p, Cas9 3 NLS, and 1.4µl reaction buffer. The mixture was incubated at room temperature for 5 minutes to form RNP complexes. 50µl protoplast was added to the mixture with 12.5µl PEG/CaCl_2_ and about 1ug DNA donor, and incubated on ice for 20 minutes. 500µl PEG/CaCl_2_ was then added and the mixture was incubated at room temperature for 5 minutes. 1 ml SCT solution was then added to the mixture and the mixture was spread on the regeneration plates. After incubation at 37°C for 20-24 hours, an over layer of 5ml regeneration medium with 600ng/ml Hygromycin B (Sigma-Aldrich) was added and put back to incubate until the colonies appeared on the regeneration plates. The colonies were transferred to 60mm small regeneration plates with 100ng/ml Hygromycin B antibiotic. The control was carried out without the RNP complex and donor DNA and replaced by 1mM Tri-HCl. An over layer with and without antibiotic Hygromycin B over layer was added to the control plates. The schematic diagram of Cas9 and sgRNA RNP complex directed with PEG-mediation transformation into *C. cinerea* to generate target gene disruption is shown in Fig. 1.

**Fig. 1.**
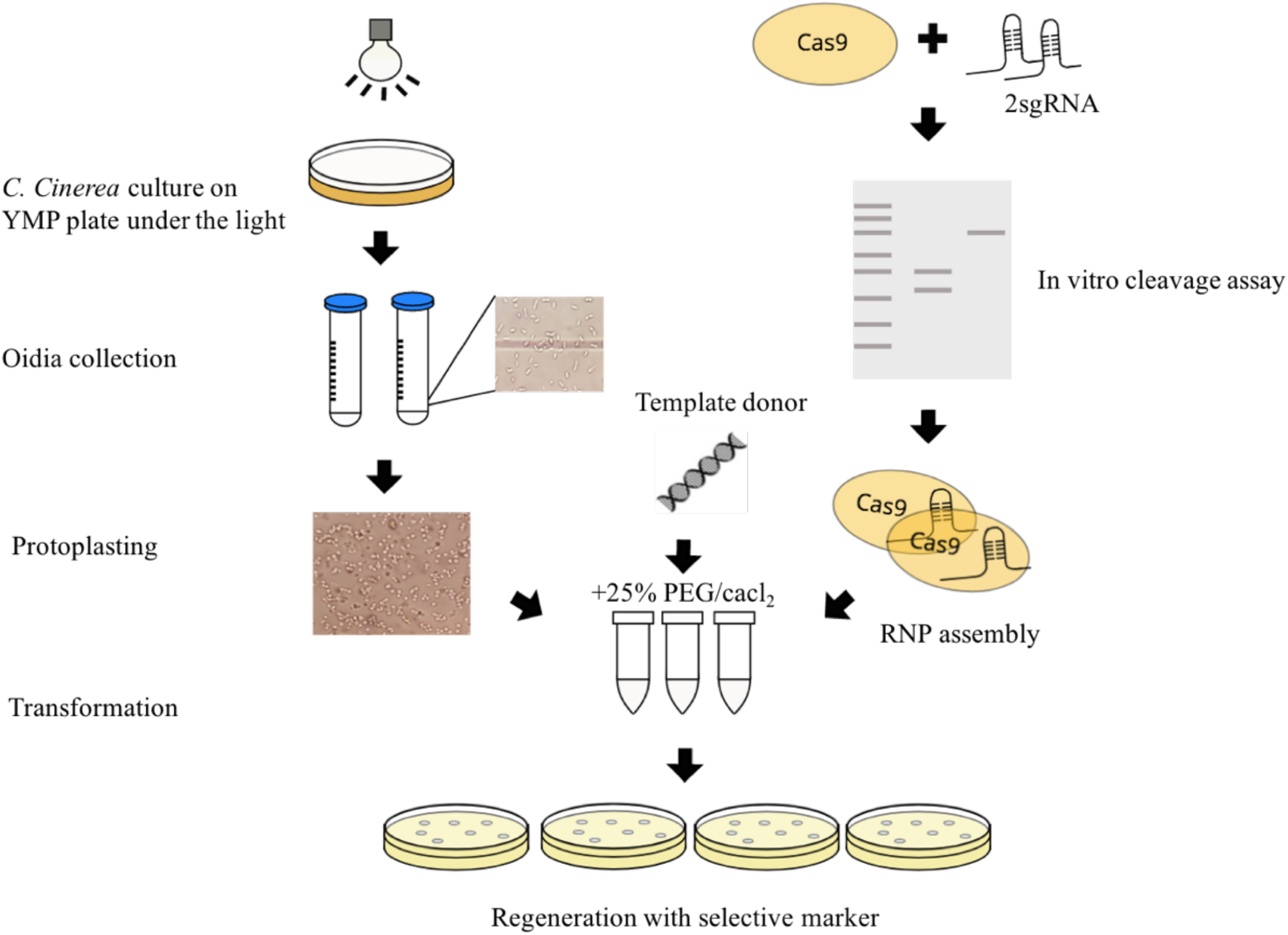
The schematic diagram of sgRNAs and Cas9 RNPs directed with PEG mediation transformation to *C. cinerea* to generate target gene disruption.

### Colony PCR

To further confirm the template donor had been correctly inserted, the minute mycelium was picked from individual colonies after 4 days of incubation at 37°C with sterilised tips and soaked in 50µl TE buffer. Using The mycelium was heat up in a microwave (600W) for 1 minute, vortexed vigorously, and put into a freezer for incubation for 30 minutes, before centrifugation for 5 minutes at 10,000rpm. 3µl supernatant, which acted as templates, was added to 7µl KAPA HiFi PCR solution and three pairs of primer (PKA1_TMP_FP + HPH_in_RP, HPH_in_FP + PKA1_TMP_RP, PKA1_TMP_FP + PKA1_TMP_RP) to determine the insertion of template donor into the genome of colony candidates. The primers for verification are listed in Table 2.

### Phenotype analysis

For assay of mycelial growth and fruiting body formation, a piece of mycelium 5 mm in diameter was cut from the edge of the growing colony of wild type strains and *CcPka1* mutant strains were inoculated on YMG plates and YMG plates with 1.5% glucose, 70mM LiCl and 500µM CHIR99021 trihydrochloride, respectively. The wild type strain was also grown on YMG plates with 100nM cAMP and incubated in the dark at 37°C for 6 days. After the mycelium had reached the edge of plates, the plates were incubated at 28°C in a 12/12 h light/dark cycle for fruiting body development. Besides, wild type and *CcPka1* mutants were inoculated in YMG plates with 1.5% glucose and incubated at 28°C in an all-dark environment.

### Glucose and glycogen assay

To measure the amount of glycogen and glucose present at the stage of *C. cinerea* development, wild-type strain and Δ*CcPka1* mutant strains were grown on YMG plates at 37°C and harvested after 4 days of incubation as well as the hyphal knots formed in Δ*CcPka1* mutants. The collected mycelia and hyphal knots were ground into powder using liquid nitrogen. The powder was separated into two sets for glucose, glycogen, and weight measurements. For glucose measurement, the amount of glucose in wild type and Δ*CcPka1* mutants was measured colourimetrically by using a dinitrosalicylic acid (DNS) reagent with 40% potassium sodium tartrate as suggested by Miller (1959). 300µl of DNS reagent was added to 150µl sample and incubated in a bath of boiling water for 5 mins. After incubation, distilled water of up to 1.5 ml was added. The absorbance was measured at 540 nm and a standard curve with standard glucose solution was plotted to calculate the glucose concentration using the DNS reagent method. For glycogen quantification, 1 ml KOH solution was added to the weighed samples and heated in boiling water in a bath for 30 mins. After heating, the sample solution was centrifuged at 14600rpm for 5mins and 100µl supernatant was added to 1.3ml 95% ethanol to precipitate glycogen. It was then centrifuged and the solution was discarded to dry the pellet. 200µl NH4Cl was added to dissolve the pellet. 130µl iodine solution was added to 100µl samples. The absorbance was measured at 495 nm and a standard curve with standard glycogen solution was plotted to calculate the glycogen concentration using iodine solution method.

### RNA extraction

Mycelium of the wild-type strain and mutants was incubated at 37°C in total darkness for 4 days and then harvested for RNA extraction. Another set of RNA extraction was carried out from the mutant plates after the formation of hyphal knots. The RNA extraction was carried out using RNeasy Plant Mini Kit (Qiagen). RNA was first treated with the TURBO DNA-free kit according to the manufacturer’s instructions. The quality and quantity of the extracted RNA were assessed on 1.5% agarose gels and a NanoDrop Lite Spectrophotometer (Thermo Fisher Scientific).

### cDNA synthesis and quantitative real-time PCR

Approximately 500 ng of total RNA was used to synthesize cDNA using the iScript gDNA Clear cDNA Synthesis Kit (Bio-Rad) according to the manufacturer’s instructions. The synthetic cDNA of each sample was used as the templates for PCR assay with specific primer in RT-PCR checking. SYBR green-based qRT-PCR was then performed using SsoAdvanced Universal SYBR Green Supermix (Bio-Rad) on Applied Biosystems 7500. The relative expression levels of the target genes were calculated using 2^^-ΔΔCT^ method. Beta-tubulin was used as an endogenous control for normalization. The primers of the genes of interest are shown in Table 3.

**Table 3.**
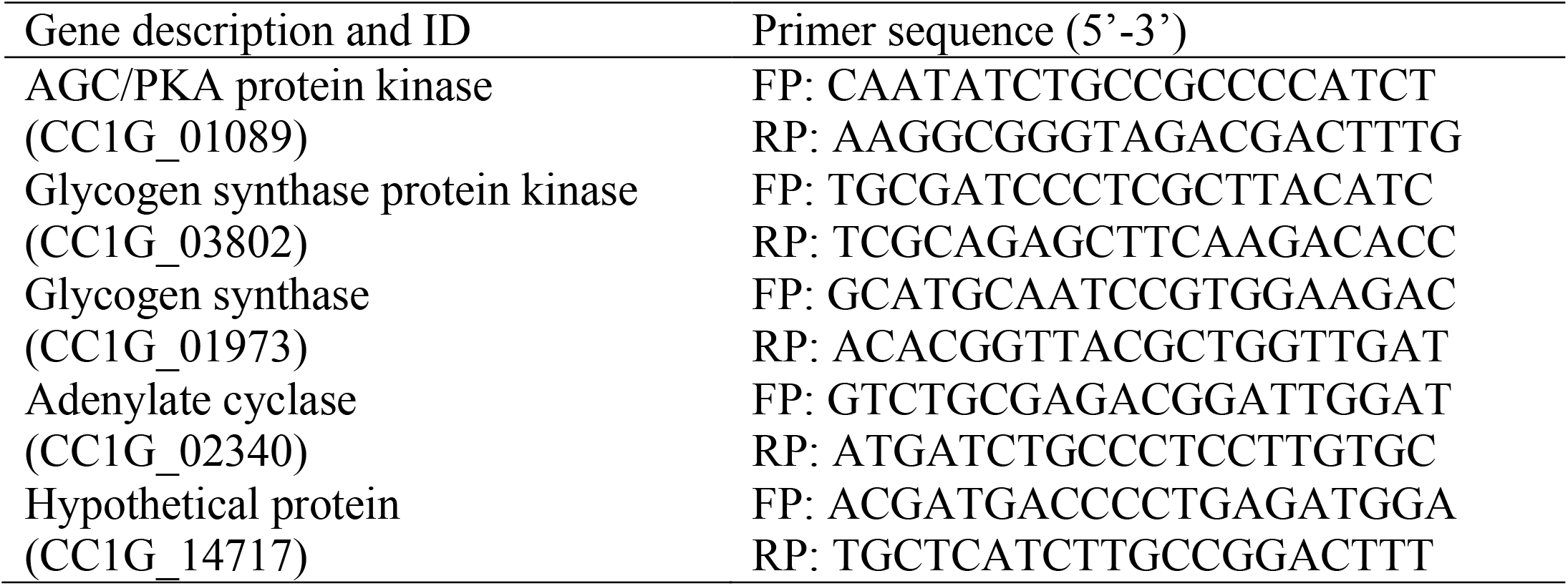
Primers for the RT-PCR and qRT-PCR of *CcPka1*gene and interested genes.

## Results

### Development of CRISPR Cas9 RNP knockout of PKA Cleavage efficiency of RNP complexes

To ensure good cleavage efficiency of the sgRNAs and Cas9 RNP complexes, we performed an *in vitro* cleavage activity assay with the target gene fragment including target cut site. After incubation for an hour, the cleavage result was visualised by 1.5% agarose gel (Fig. 2). SgRNA1 and SgRNA2 with Cas9 RNP complex successfully cut 88% and 91% of the target gene template, respectively. Meanwhile, almost all the target gene template was cleaved when two RNP complexes were combined. The result indicated that one RNP complexes and two RNP complexes combined both have high cleavage efficiency.

**Fig. 2.**
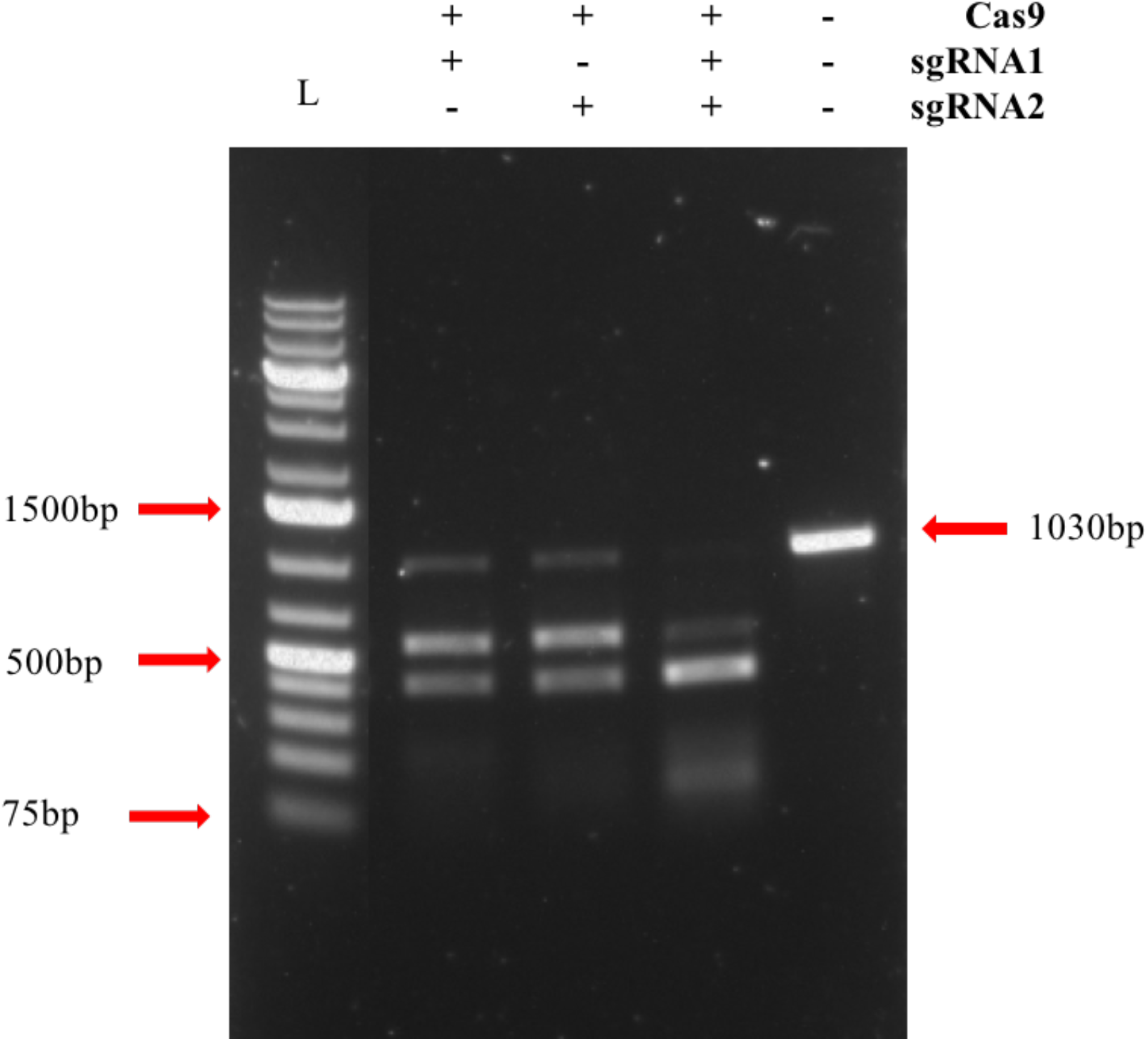
*In vitro* cleavage activity of sgRNAs and Cas9 RNP complexes by gel electrophoresis. About 1030bp target gene template was mixed with one sgRNA and Cas9 complex or two sgRNA and Cas9 complex, and incubated at 37°C for 1 hour. L: 1kb ladder.

### Target disruption of PKA in *C. cinerea* using a selective marker

To knock out PKA, a target gene without any observable phenotype, we performed a target gene disruption by using two sgRNA and Cas9 RNP complexes with a template donor. The template donor included hygromycin resistant gene. Around 4 days after two transformations, 40 colonies and 61 colonies grew on regeneration plates with the antibiotic Hygromycin B (600ng/ml). They were grown on regeneration plates with Hygromycin B (100ng/ml). Ten transformants were randomly chosen and verified by PCR with specific primer to the template donor inserted into the target cut site. The amplicons were visualised by 1.5% agarose gel electrophoresis. (Figs. 3A&B). Half of the transformants had the template donor successfully inserted in the target cut site. However, the sizes of some amplicons size were larger than the expected size. The amplicons were purified and sequenced. The template donor was inserted in one cut site.

**Fig. 3.**
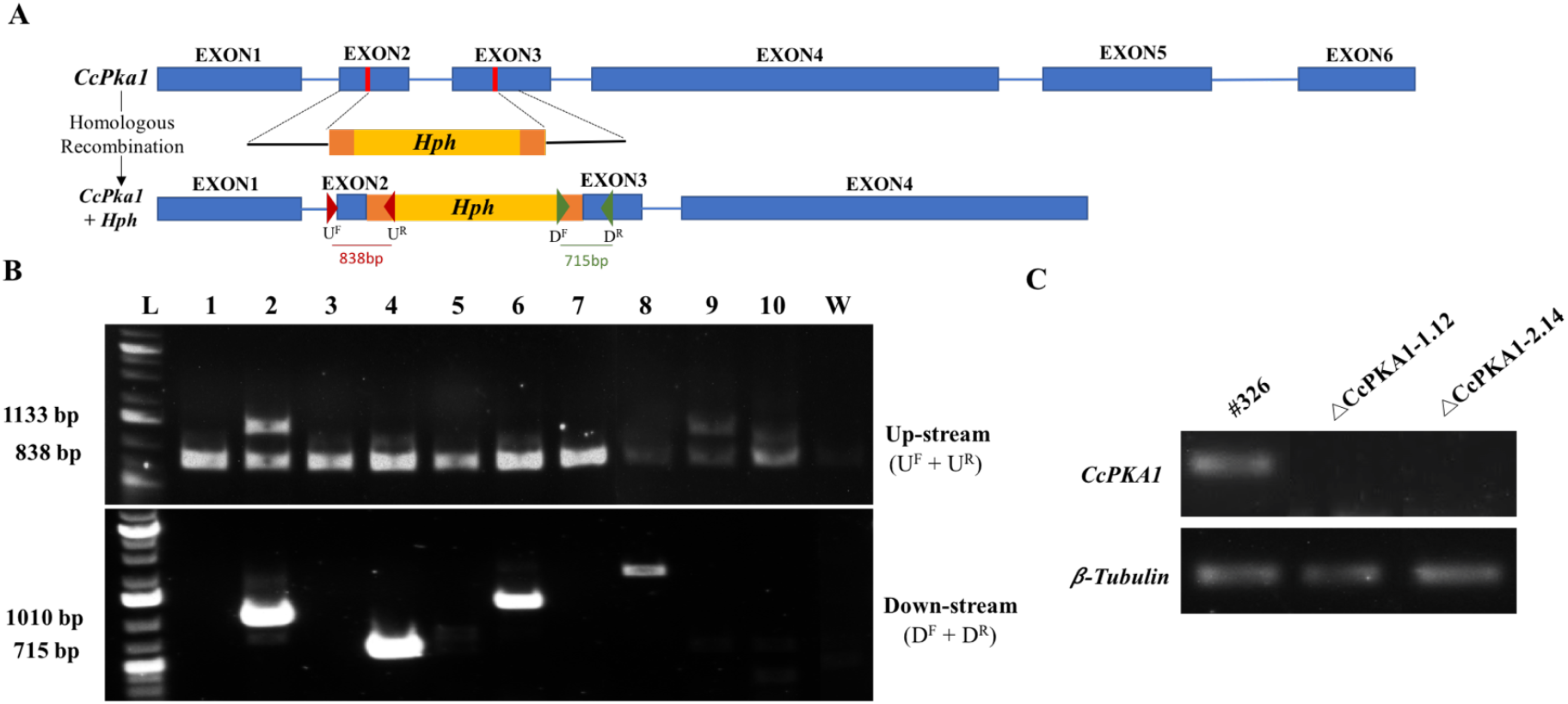
Verification of *CcPka1* mutants generated by Cas9 RNP transformation. A. Gene disruption by homologous recombination to insertion site with hygromycin cassette; B. Verification of the insertion of template donor into genome of Δ*CcPka1* mutants by PCR, L: 1kb ladder, W: wild type #326, 1-10: putative Δ*CcPka1* mutants. UP-stream PCR product amplified with primers PKA1_TMP_FP + HPH_in_RP, Down-stream PCR product amplified with HPH_in_FP + PKA1_TMP_RP; C. The expression of *CcPka1* in wild-type and two Δ*CcPka1* mutant strains by RT-PCR. The expression of Beta-tubulin acts as an internal control.

To confirm the disruption of the*CcPka1*gene, its expressions in selected mutants were tested by RT-PCR. There was no expression of *CcPka1*gene in these selected mutants when compared with the wild-type strain. (Fig. 3C)

### PKA mutant strains grew more slowly than wild-type strain

To investigate the role of PKA in the cAMP-dependent signaling pathway in mycelial growth, the Δ*CcPka1* mutants and wild type strains were cultured on YMG plates for 6 days. The Δ*CcPka1* mutants grew more slowly than the wild type strains (Fig. 4). The growth rate differences of Δ*CcPka1* mutants were greater than that of wild type strains after 5 days of incubation. Δ*CcPka1* mutants required one more day to reach the edge of the plate. The results indicated that PKA mutation affected the mycelial growth of *C. cinerea*.

**Fig. 4.**
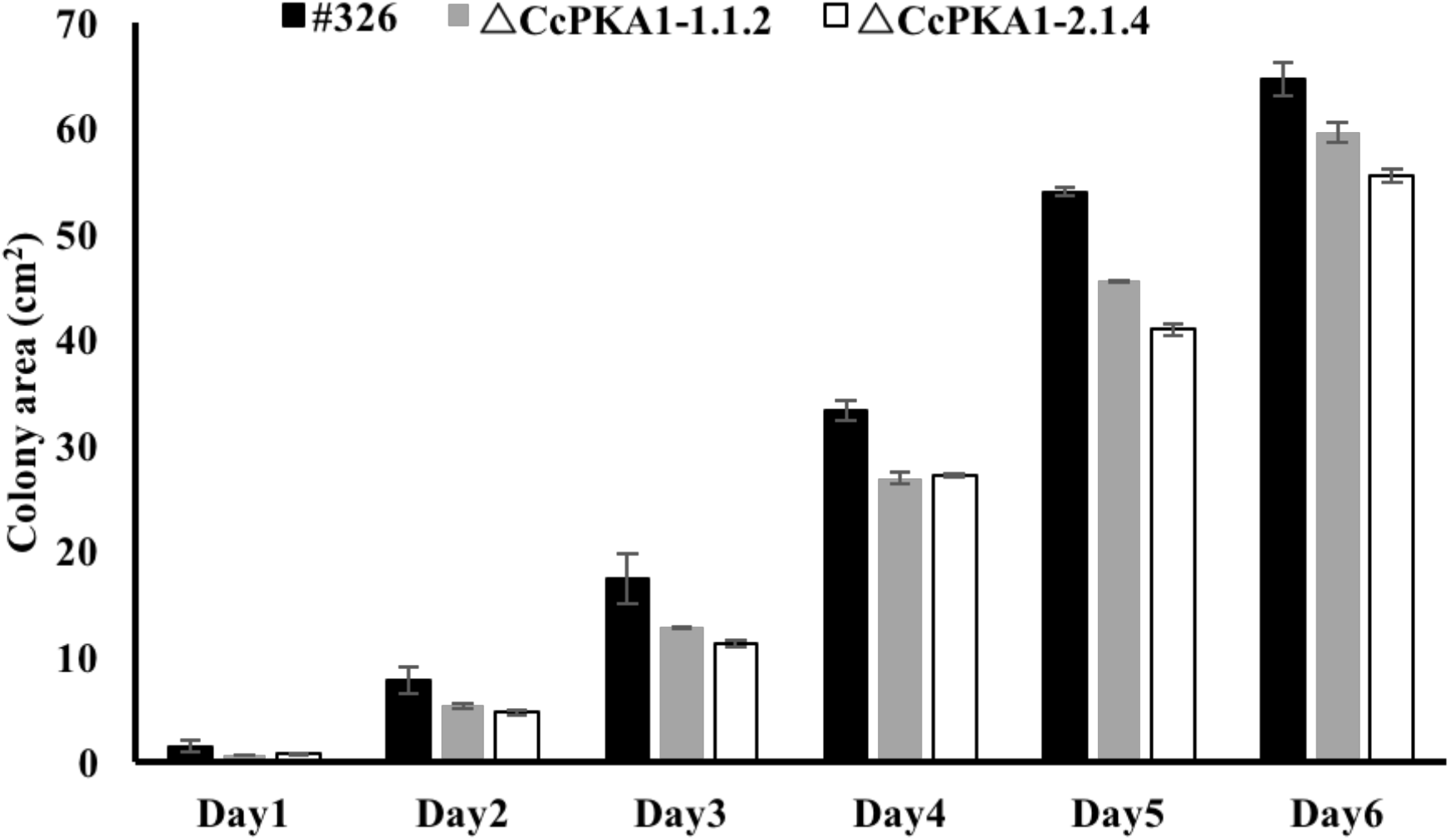
The mycelium growth of wild-type strain and Δ*CcPka1* mutants. The colony areas of wild type and two Δ*CcPka1* mutants were measured for 6 days. Each experiment had biological triplicates. Statistical analysis was preformed using one way pair sample of mean of Student’s t test when compared with the wild-types strains.

### PKA mediated the intracellular glucose and glycogen level

To determine the role of PKA in *C. cinerea*, wild-type and Δ*CcPka1* mutants were grown on YMG plates for 4 days to observe the time required to form hyphal knots. Intracellular glycogen and glucose levels were measured in Δ*CcPka1* mutants and wild type strains (Fig. 5). The intracellular glycogen level in Δ*CcPka1* mutants was reduced compared to that of the wild-type strain. After transferring the wild-type and Δ*CcPka1* mutants cultures to a 12 h/12h light/dark cycle at 28°C, the glycogen level in the wild-type strain decreased. Meanwhile, the intracellular glycogen level in Δ*CcPka1* mutants was significantly lower than that of the wild-type strain. For the glucose level, the mycelium of Δ*CcPka1* mutants had significantly higher intracellular glucose content than the wild-type strain. The glucose contents in the wild-type mycelium were similar at two time points. Despite the decreased glucose contents of Δ*CcPka1* mutants after the formation of hyphal knots, the intercellular glucose levels were still higher than that in the wild-type strain. The results suggested that accumulated intracellular glucose might enhance the fruiting body development of *C. cinerea*.

**Fig. 5.**
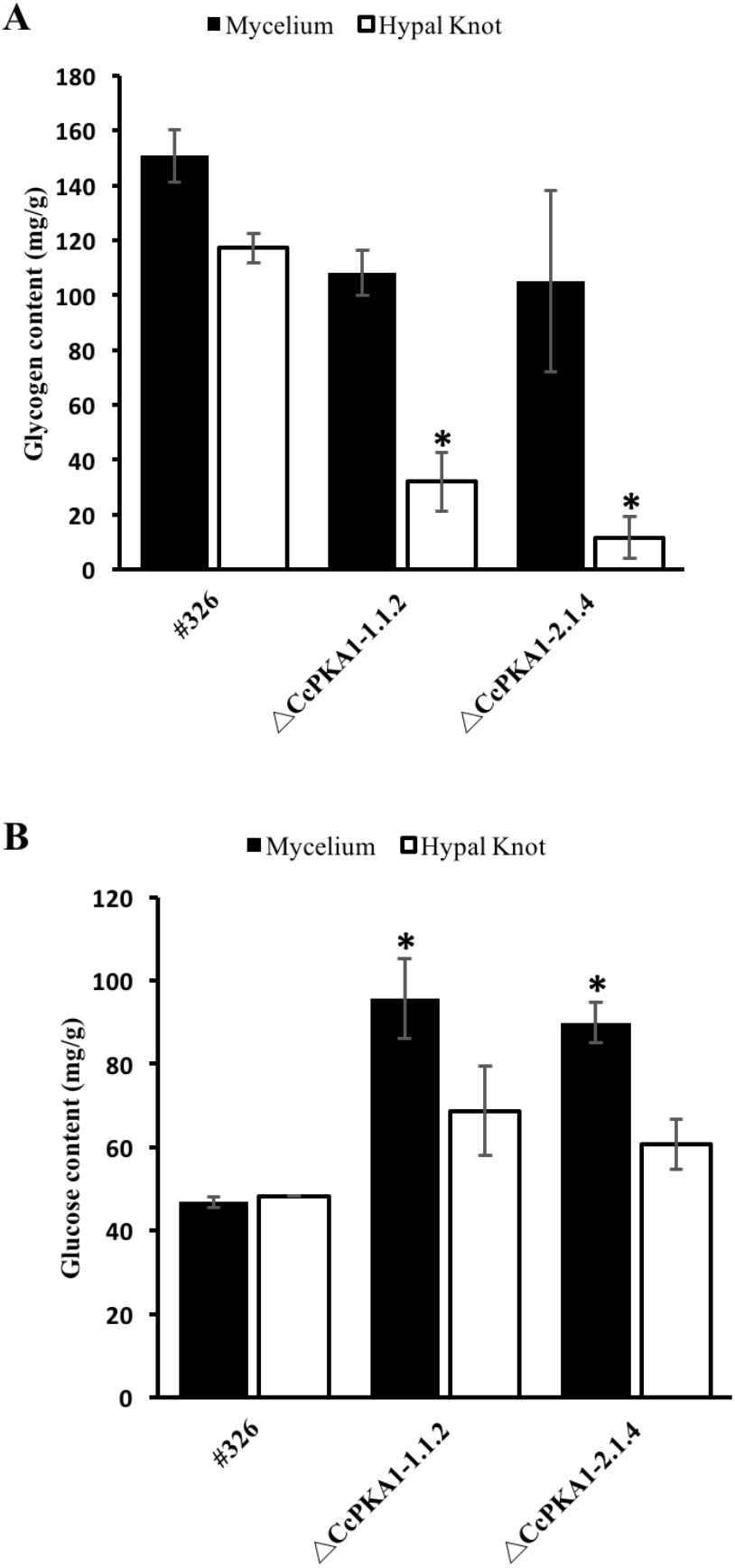
The intracellular glycogen and glucose levels in the wild-type strain and *CcPka1*mutant strains. A. The intracellular glycogen content in the wild type and 2 Δ*CcPka1* mutants. B. The intracellular glucose content in the wild type and 2 Δ*CcPka1* mutants. Each experiment was done in triplicate. Statistical analysis was preformed using one way pair sample of mean of Student’s t test when compared with the wild-types strains; *: P<0.05.

### The role of PKA in fruiting body development

*C. cinerea* is a light-dependent and nutrient depleting mushroom-forming fungi (Kües, 2000). To investigate the function of PKA in the fruiting body development of *C. cinerea*, we cultured the wild-type and Δ*CcPka1* mutants in regular YMG plates. Though the Δ*CcPka1* mutants grew more slowly than the wild type, their hyphal knots were formed one to two days earlier than those of the wild type and developed into fruiting body. PKA might have played an essential role in glucose signaling when wild type and Δ*CcPka1* mutants were grown on YMG plate with 1.5% glucose. The high glucose content would be a high nutrient condition. The mycelial growth of all strains was almost the same but grew slower in high glucose content YMG plates when compared to regular YMG plates. After mycelia reached the edge of plates, the plates were put under a 12h/ 12h light-dark cycle. The Δ*CcPka1* mutants prompted the fruiting initiation under high glucose condition (Fig. 6). The fructification of mutant strains initiated after 4 days of incubation. In contrast, the wild-type strain remained in the mycelium stage. When the mutants developed into mature fruiting body, wild-type strain only started to form primordium. These results suggested that PKA could be a nutrient signal transductor and could negatively regulate the fruiting body development of *C. cinerea*.

**Fig. 6.**
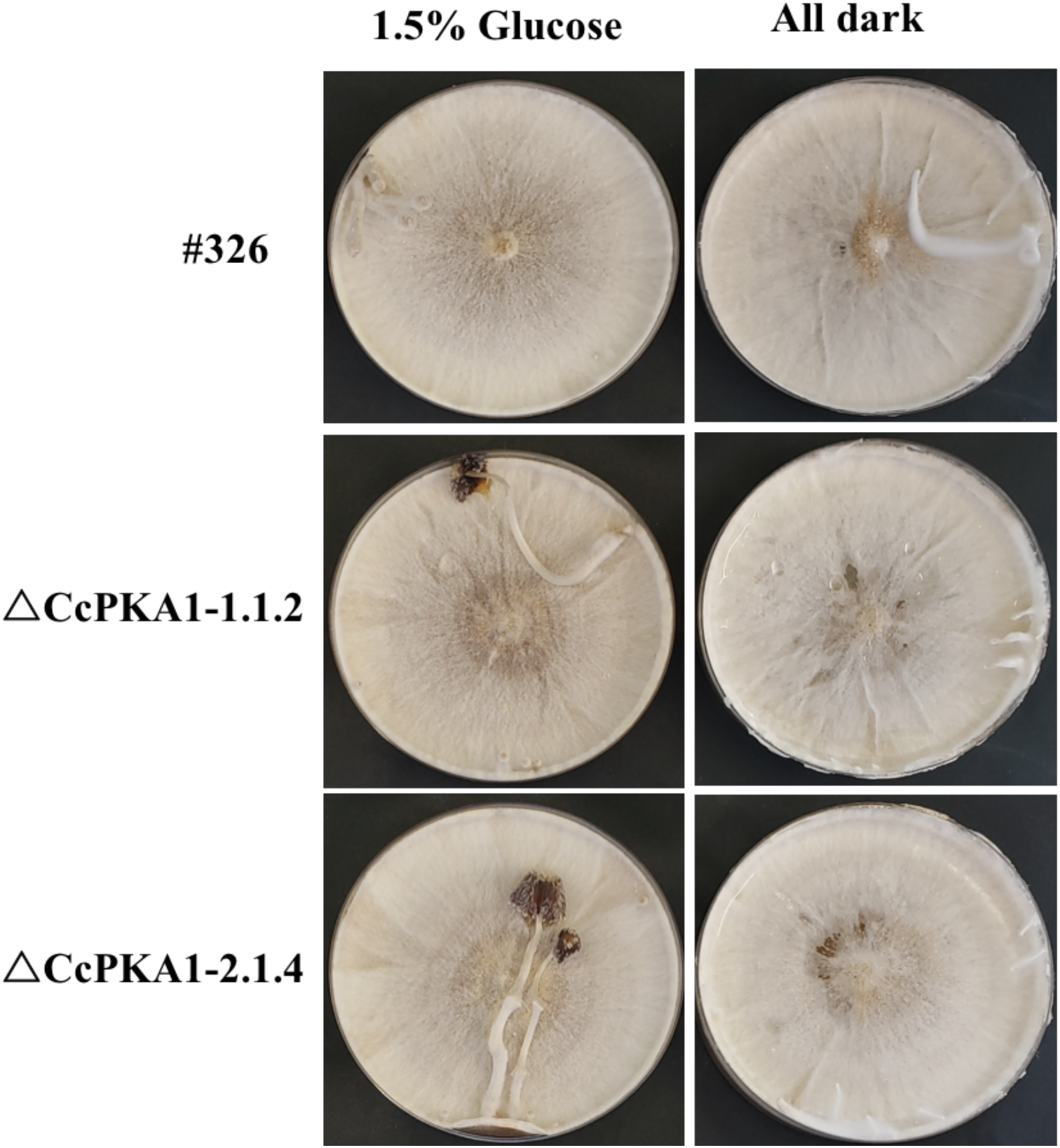
The fruiting body development of wild-type and *CcPka1*mutants under different extreme growth conditions. When cultured in high glucose YMG plates, the Δ*CcPka1* mutants formed fruiting body much earlier than the wild-type strain. The fruiting body of Δ*CcPka1* mutants autolyzed while the wild type just formed young fruiting body. Neither strains could form normal fruiting body in the all-dark environment.

**Fig. 7.**
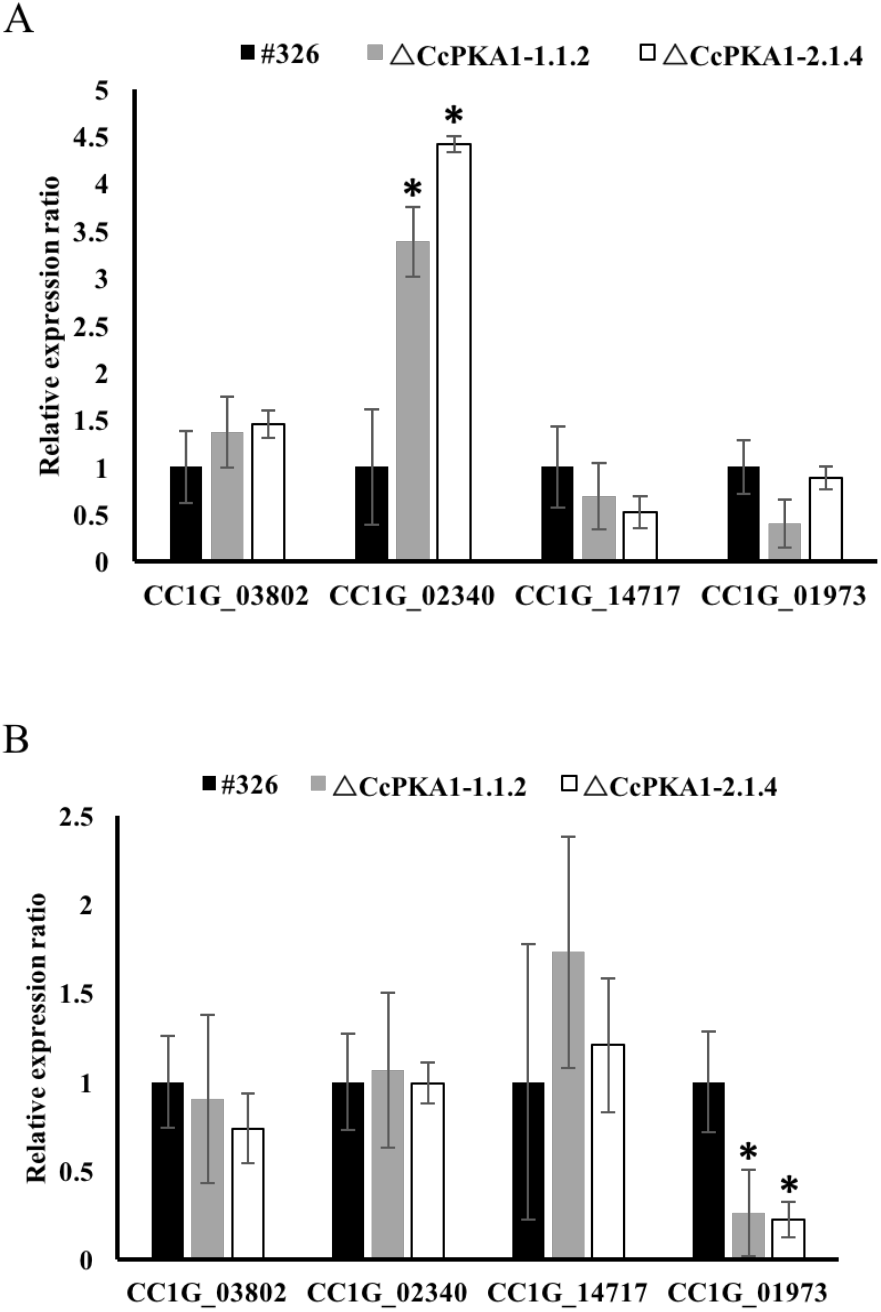
The gene expression of interested genes in wild-type strain and Δ*CcPka1* mutant strains. A. The expression of the genes of interest in wild-type and 2 Δ*CcPka1* mutant strain during the mycelium stage; B. The expression of the genes interest in wild type and 2 Δ*CcPka1* mutant strain during the formation of hyphal knots. The genes of interest were GSK3 (CC1G_03802), adenylyl cyclase (CC1G_02340), ras GTPase activator (CC1G_14717), and glycogen synthase (CC1G_01973). Each experiment was carried out in triplicate. Statistical analysis was preformed using one way pair sample of mean of Student’s t test when compared to the wild-type strains; *: P<0.05.

### Deletion of *CcPka1* affected the gene expression

In cAMP-dependent PKA pathway, adenylyl cyclase is the upstream target of PKA to synthesize cAMP which regulates the activity of PAK (Rolland et al. 2000). Therefore, we selected two upstream components, CC1G_02340 and CC1G_14717, in this signaling pathway and evaluated their expressions. CC1G_02340 gene encodes adenylyl cyclase and CC1G_14717 gene, which is a hypothetical protein, encodes an ortholog of *Lentinula edodes* ras GTPase activator with 63% homology. During the mycelium stage, the relative expression of adenylyl cyclase (CC1G_02340) was upregulated 3 to 4-fold in the *CcPka1*mutant strain compared to the wild-type strain. We speculated that the Δ*CcPka1* mutants, with no functioning PKA, transmitted a low glucose signal that induced the expression of adenylyl cyclase to synthesize cAMP. To modulate cAMP level during hyphal knot induction, the gene expression of adenylyl cyclase (CC1G_02340) was not significantly changed when compared to the wild-type strain. Compared to the wild-type strain, the gene expression of ras GTPase activator (CC1G_02340) was reduced during mycelium stage and then increased when hyphal knots were formed. The relative expression of glycogen synthase (CC1G_01973) was downregulated in Δ*CcPka1* mutants compared to the wild-type strain in the mycelium stage as well as when the hyphal knots were formed, so that glycogen content was reduced severely. In contrast, the expression of glycogen synthase protein kinase (CC1G_03802) increased when compared to the wild-type strain in the mycelium stage. However, its expression reduced slightly during the formation of hyphal knots. These results indicated that PKA could affect the expression of glycogen synthase and PKA could not be only one protein kinase affect the expression of glycogen synthase kinase.

### GSK3 could be downstream of cAMP-PKA pathway

In mammalian studies, the activation of glycogen synthase kinase was inhibited by the phosphorylation mediated by protein kinase A (Fang et al. 2000). To investigate the relationship between PKA and GSK3 in *C. cinerea*, we applied 500µM CHIR99021 trihydrochloride, a specific GSK3 inhibitor, and 70mM LiCl to wild-type strain and Δ*CcPka1* mutant strains during incubation (Fig. 8). After two weeks of incubation, both strains remained in the mycelium stage and did not fruit. The results showed that GSK3 was downstream of PKA in the signaling pathway and suggested that GSK3 might be a target of PKA. No the other hands, the results were possible as GSK3 and PKA were independent regulators in the control of fruiting.

**Fig. 8.**
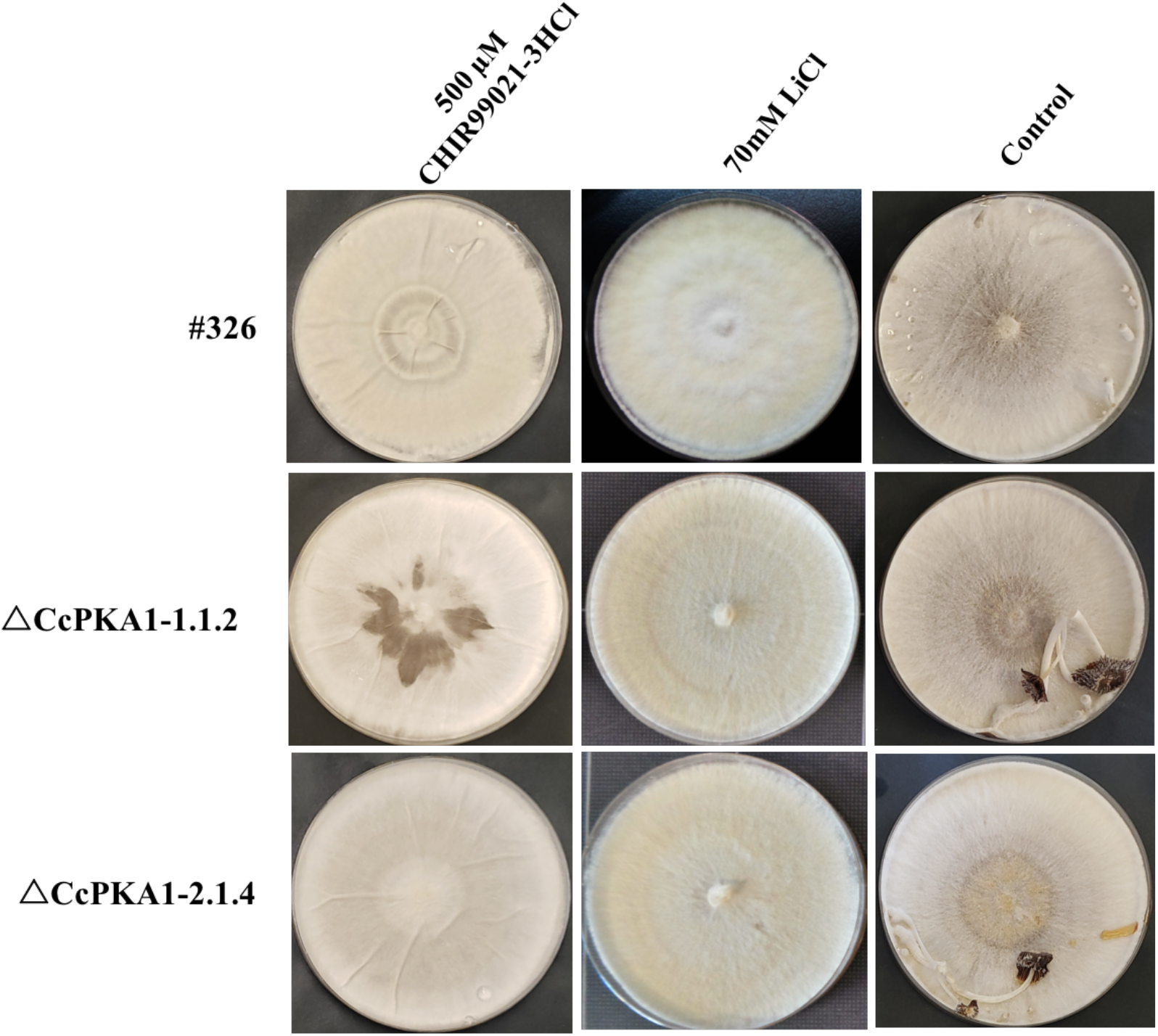
The inhibitor intervention in fruiting body development of wild-type and *CcPka1*mutants. The adding of two GSK3 inhibitors to wild type and Δ*CcPka1* mutants interrupted the fruiting body formation in both strains after incubation for two weeks. Control groups were without addition of GSK3 inhibitors.

## Discussion

### Advantages of CRISPR-Cas9 RNP

Target gene manipulations, such as gene knockouts, are important molecular methods to understand the functions of genes in various processes during fungal development, fruiting body morphogenesis, metabolic regulation, and meiosis (Kamada 2002; Kues 2000). Especially, a better understanding of gene functions in sexual development would allow us to improve fungal strains by appropriate genetic modification. However, due to the low homologous recombination frequency of gene targeting in *C. cinerea*, there are only a few studies that report the functional analysis of target genes by transforming a selectable marker flanked with homologous sequence (Binninger et al. 1991; Nakazawa et al. 2011).This not only happens in *C. cinerea* but also in *Schizphyllum commune* and *Pleurotus ostreatus* (Ohm et al. 2010; Salame et al. 2012). Only one to three transformants with a target gene deletion have been obtained in previous studies. Some studies have attempted to improve the homologous recombination(HR) efficiency by disruption of Ku80/Ku70 to reduce the incidence of non-homologous end joining (NHEJ), which is a pathway competing with HR (De Jong et al. 2010; Ninomiya et al. 2004; Salame et al. 2013). However, the lack of *Ku80/Ku70* has some drawbacks, such as reducing the protoplast regeneration rate and influencing sexual development (De Jong et al. 2010; Nakazawa et al. 2011). Herein, a simple and effective gene manipulation method, the CRISPR-Cas9 system with RNP in *C. cinerea* combining with two sgRNAs, was developed. About 50% of transformants were successfully integrated template donor into *CcPka1* gene, while only 8% of transformants occurred in the traditional mutagenesis methods. In other mushroom-forming fungi, the RNP with template donor, including the appropriate flanking region, also enhanced the success rates of homologous gene replacement (Vonk et al. 2019). With this approach, two or more target genes modified with different sgRNAs could be achieved together with the same or different selective markers to study the gene functions in mushroom-forming fungi.

## The role of cAMP in cAMP-PKA signaling pathway

Fruiting body development of basidiomycetes requires several environmental conditions including appropriate nutrient, light, temperature and humidity. Changes in these conditions would induce the relevant sensors to receive and pass the signal to signal transduction pathways (Lengeler et al. 2000). Signal transduction pathways have specific components that occur in a particular condition. The role of nutrient-related signaling pathways, mainly cAMP-dependent PKA pathway, in the growth and development regulation and carbohydrate metabolism in filamentous fungi and mushroom-forming fungi, has been studied (Bruno et al. 1996; Li and Borkovich 2006; Yamagishi et al. 2005). In this study, we investigated the role of PKA pathway in the fruiting body development of *C. cinerea* under unfavourable nutrient condition.

In *Schoizophyllum commune*, overexpression of *ScPKAC1* and *ScPKAC2*, which encoded cAMP-dependent protein kinase catalytic subunit, suppressed the aerial mycelium formation and fruiting body formation without affecting the intracellular cAMP level (Yamagishi et al. 2005). In contrast, overexpression of *ScGP-A*, which encoded heterotrimeric G-protein alpha in *S. commune*, also suppressed aerial mycelium formation and increased the intracellular cAMP level (Yamagishi et al. 2005). In this study, the Δ*CcPka1* mutants reduced mycelial growth and induced fruiting body. The results confirmed that the inactivation of PKA could be responsible for inducing the fruiting body development without suppression of its target.

In Δ*CcPka1* mutants, the expression of adenylyl cyclase gene (CC1G_02340) increased greatly, suggesting that the cAMP level in the cytoplasm might increase. The increased cAMP level could be caused by the active G-protein alpha. The G-protein-couple receptor (GPCR) system takes up glucose from growth medium and activates cAMP synthesis (Rolland et al. 2000). cAMP is crucial for cell growth and differentiation in fungi (Kues 2000). Fruiting body formation of *C. macrorhizus* is promoted by the addition of cAMP (Uno and Ishikawa 1973). In this study, adding exogenous cAMP to *C. cinerea* reduced the mycelial growth rate and stimulated fruiting body formation. The amount of intracellular glucose in the exogenous cAMP-treated wild type strain was more than that in the wild-type strain without exogenous cAMP. In *Cryptococcus neoformans* and *Ustilago maydis*, G-protein alpha responds to the signal of nutrient deprivation that induces cAMP generation to activate cAMP-dependent PKA (Alspaugh et al. 2002; Kaffarnik et al. 2003). In contrast, Cheng et al. and Yamagishi et al. studies showed that the expression of PKA was reduced by half in *C. cinerea* between mycelium and primordium in *S. commune* as well when the expression of adenylyl cyclase remained unchanged (Cheng et al. 2013; Yamagishi et al. 2005). We speculated that when cAMP level increased, cAMP might bind to PKA and release the suppression of PKA to its downstream target other than activating PKA.

## Relationship between GSK3 and PKA

In *N. crassa*, the mutation of *cr-1* gene, which encoded adenylyl cyclase, increased glycogen levels during vegetative growth (Freitas et al. 2010). Similarly, Thevelein and Winde (1999) showed that increased cAMP level decreased glycogen level and repressed gluconeogenesis (Thevelein and Winde 1999). These studies indicated the role of cAMP-PKA pathway in the regulation of intracellular glycogen level. We speculated that PKA might serve as a negative regulator to control the glycogen reserves. Furthermore, the glycogen mobilization might be regulated by PKA pathway directly or indirectly (Freitas et al. 2010). In this study, the intracellular glycogen content was reduced greatly in Δ*CcPka1* mutant strains during the formation of hyphal knots, the initial step of fruiting body formation. Glycogen synthase kinase (GSK3), a Ser/Thr protein kinase, phosphorylated and inactivated glycogen synthase (Osolodkin et al. 2011). In *Fusarium graminearum*, glycogen content increased in a Δ*fgk3* mutant, showing that GSK3 inhibited glycogen synthase (Qin et al. 2015). We suggested that active GSK3 also inhibited glycogen synthase activity in *C. cinerea*. Zhou et al., (2017) reported that GSK3 was regulated transcriptionally by a MAP kinase in *Magnaporthe oryzae* (Zhou et al. 2017). Apart from the MAP kinase regulation, mammalian cAMP-dependent PKA inhibited the activity of GSK3 in response to changes in intracellular cAMP level (Fang et al. 2000). The interaction between PKA and GSK3 is still unclear. GSK3 inhibitor intervention showed that Δ*CcPka1* mutants remained in the vegetative stage and could not form fruiting body, indicating that GSK3 in *C. cinerea* might be downstream of PKA, that PKA might suppress the activity of GSK3 during the mycelium stage, and that the repression of PKA to GSK3 was reduced during the fruiting body development.

## Conclusion

Using the CRISPR/Cas9 system, the knockout of *CcPka1* in *C. cinerea* was successfully carried out with an almost 50% success rate. The Cas9 ribonucleoprotein and the selective maker donor made the gene editing process more convenient. Therefore, CRISPR/Cas9 RNP could advance the gene mutagenesis of the genes of interest in fungal study.

Meanwhile, the Δ*CcPka1* mutants grew slowly in the mycelium stage, but they formed fruiting body more quickly than the wild type. The results showed that PKA negatively regulated the fruiting body development and transmitted the nutrient signal to mediate fruiting body development of *C. cinerea*. Since PKA mediates intracellular glucose and glycogen levels, accumulated intracellular glucose might enhance the fruiting body development of *C. cinerea*. Moreover, the increased cAMP might bind to PKA, then the suppression of PKA might release to its downstream target, other than activating PKA during mushroom forming induction. From GSK3 inhibitor intervention, the result showed that GSK3 might be the downstream target of PKA. Yet, GSK3 and PKA are independent regulators in the control of fruiting. Further study to investigate the interaction between PKA and GSK3 in *C. cinerea* is required.

## Acknowledgment

This study was supported by the RGC General Research Fund (GRF14107514 and GRF 14103817) from the Research Grants Council of the Hong Kong SAR.

## Notes

### Competing Interest Statement

The authors have declared no competing interest.

